# Integrated Stress Response Inhibition Provides Sex-Dependent Protection Against Noise-Induced Cochlear Synaptopathy

**DOI:** 10.1101/2020.06.17.158303

**Authors:** Stephanie L. Rouse, Ian R. Matthews, Jiang Li, Elliott H. Sherr, Dylan K. Chan

## Abstract

Noise-induced hearing loss (NIHL) is a common health concern with significant social, psychological, and cognitive implications. Moderate levels of acoustic overstimulation associated with tinnitus and impaired speech perception cause cochlear synaptopathy, characterized physiologically by reduction in wave I of the suprathreshold auditory brainstem response (ABR) and reduced number of synapses between sensory hair cells and auditory neurons. The unfolded protein response (UPR), an endoplasmic reticulum stress response pathway, has been implicated in the pathogenesis and treatment of NIHL as well as neurodegeneration and synaptic damage in the brain. In this study, we used the small molecule UPR modulator ISRIB (Integrated Stress Response InhiBitor) to treat noise-induced cochlear synaptopathy in a mouse model. Mice pretreated with ISRIB prior to noise-exposure were protected against noise-induced synapse loss. Male, but not female, mice also exhibited ISRIB-mediated protection against noise-induced suprathreshold ABR wave-I amplitude reduction. Female mice had higher baseline wave-I amplitudes but greater sensitivity to noise-induced wave-I reduction. Our results suggest that the UPR is implicated in noise-induced cochlear synaptopathy, and can be targeted for treatment.

## Introduction

Noise induced hearing loss (NIHL) affects nearly 20% of adults, with significant impact on social, psychological and cognitive function. In cochlear synaptopathy, or “hidden hearing loss”, moderate noise exposure that does not cause permanent shifts in hearing thresholds can nonetheless result in permanent loss of synapses between cochlear inner hair cells (IHCs) and spiral ganglion neurons (SGNs), associated with permanent reduced amplitude of wave I of the suprathreshold auditory brainstem response (ABR)^1^. These anatomic and physiologic findings are thought to correlate behaviorally with tinnitus and impaired speech perception in complex acoustic environments, both hallmarks of human auditory functional disability^2^.

The mechanism of synaptic damage in response to noise is unknown. Recent work by our group has implicated the unfolded protein response (UPR) pathway in permanent NIHL. The UPR is a series of highly conserved processes related to endoplasmic reticulum (ER) stress that regulate protein folding and processing, and mediate downstream homeostatic and apoptotic responses. Multiple small-molecule modulators of the UPR have been developed, including ISRIB (Integrated Stress Response InhiBitor), a drug that activates eIF2B, downregulates ATF4, and leads to decreased synthesis of the pro-apoptotic UPR protein CHOP (C/EBP-homologous protein)^3^. We found that 106 dB SPL noise exposure, which produces permanent shifts in hearing thresholds, induces CHOP expression, and that ISRIB treatment provides partial protection against hearing loss and hair-cell death in this model^4^. These results warranted further study of the protective effect of ISRIB at lower levels of noise exposure, such as those involved in cochlear synaptopathy.

Further rationale for this approach is provided by a growing body of evidence implicating the UPR and ER stress in synaptic plasticity and neurodegeneration. UPR upregulation has been found in brains of multiple human and mouse models of neurodegeneration^5,6^. UPR dysregulation results in translational failure triggering neuronal death as well as disruption of calcium signaling, essential for maintenance of neurons. Manipulation of the UPR protects neurons in multiple neurodegenerative disease models; as such, it has been an appealing target of study for potential treatment of neurodegeneration^7^. The UPR also appears to be involved in the synthesis of proteins involved in synaptic function and structural maintenance. It is a major driver of synaptic loss through reduction in global protein synthesis rates, leading to reduction in synaptic plasticity in several neurologic diseases. Specifically, downregulation of ATF4 has been shown to promote neuroprotection, synaptic plasticity, long-term potentiation, and memory formation, suggesting that ISRIB would be effective in treatment of models of synaptic dysfunction in the central nervous system^8,9^. Indeed, ISRIB has been shown to rescue synaptic plasticity in a mouse model of Down syndrome, restore cell-specific synaptic function in a mouse model of repetitive traumatic brain injury, reverse trauma-induced hippocampal-dependent learning and memory deficits, and improve memory in wild-type and memory-impaired mice^3,10–12^.

The involvement of the UPR and efficacy of ISRIB in both neurodegeneration and synaptic maintenance in the brain led us to hypothesize similar roles for the UPR and ISRIB in the cochlea. In this study, we extend our finding that ISRIB can prevent NIHL^4^ and demonstrate that ISRIB treatment also protects against noise-induced cochlear synaptopathy.

## Results

### ISRIB protects against noise-induced cochlear synaptopathy

6-8 week-old female and male wild-type CBA/J mice were treated with a single intraperitoneal dose of 2.5 mg/kg ISRIB or vehicle 2 hours prior to exposure to 2-hour, 97.8 dB SPL, 8-16 kHz octave-band noise. ABRs were measured at post-noise-exposure days (PNE) 1, 7, 14, and 21, and temporal bones harvested. We quantified synaptic ribbons between cochlear IHCs and SGNs by identifying colocalization of signal corresponding to antibodies against CtBP2, a pre-synaptic marker, and GluA2, which labels post-synaptic glutamate receptors, in male and female mice. Consistent with previous reports^1,13^, we observed significant loss of synapses in noise-treated animals (p = 0.0003) without hair-cell loss (**Figure 1**). ISRIB protected significantly against noise-induced synapse loss compared to vehicle (p = 0.0015). No significant difference was seen in synapse numbers between ISRIB-treated mice exposed to noise and control, unexposed mice (p = 0.11). Analysis of unpaired, orphan synapses, seen as presence of a presynaptic CtBP2 puncta with no co-localized post-synaptic glutamate receptor, which occur in response to noise trauma,^13^ corroborated these results, with noise-exposed, vehicle-treated mice having more orphan synapses compared to unexposed as well as to noise-exposed, ISRIB-treated mice.

**Figure 1.**
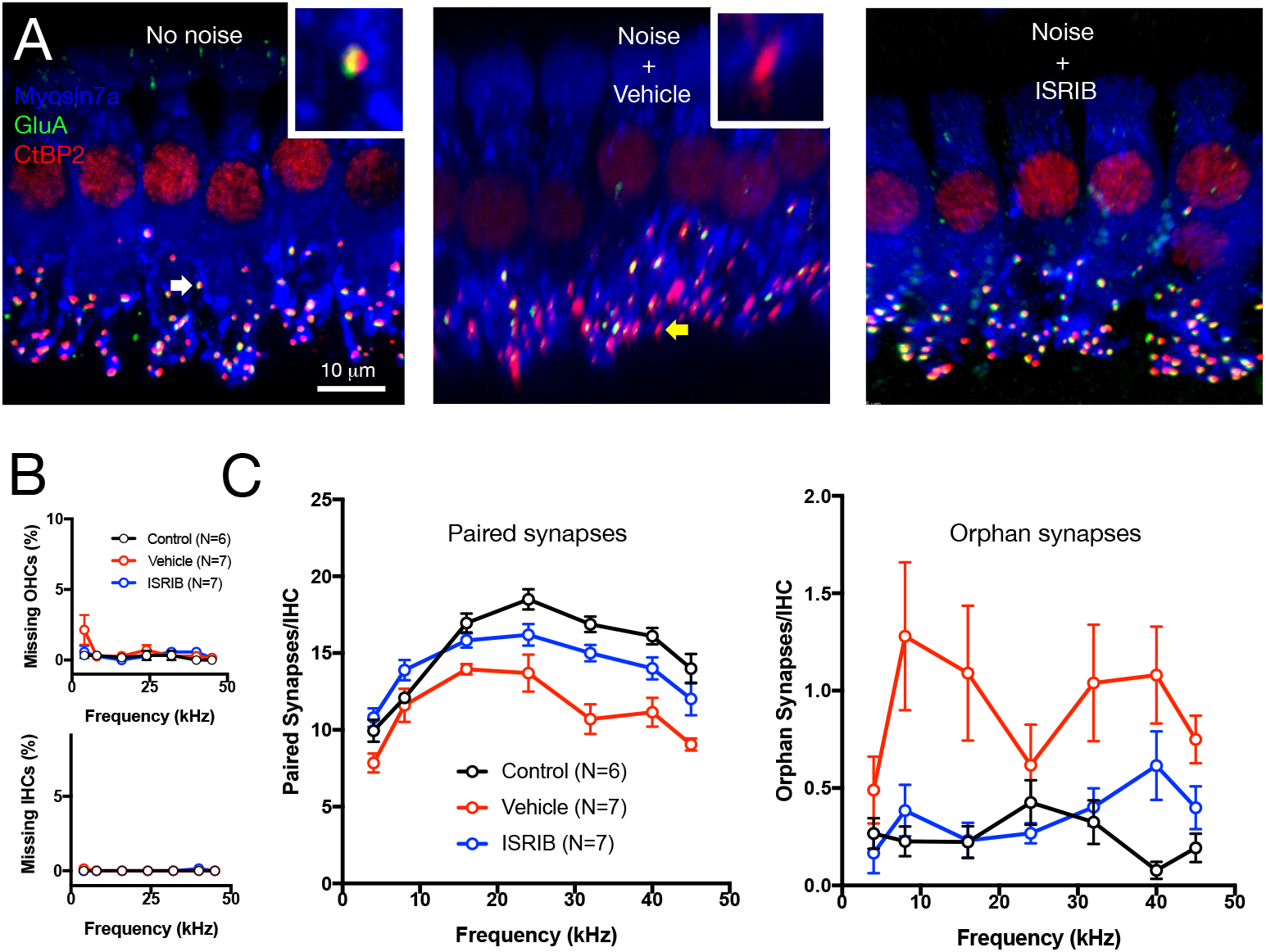
IHC-spiral ganglion neuron synapse count after noise exposure and ISRIB treatment. Female and male CBA/J mice were exposed to 2-hour, 97.8 dB 8-16 kHz octave-band noise after treatment with ISRIB or vehicle. Control mice were not exposed to noise. 45 days after noise exposure, cochlear synapses were labelled with CtBP2 and GluA2 and quantified at locations corresponding to the indicated frequencies. **A.** Representative images from the 24-kHz location are shown for mice under the indicated conditions. Arrows highlight intact synapses (white; left inset) and orphan ribbons (yellow, middle inset). Scale bar = 10 mm. **B.** Hair-cell counts demonstrated minimal-to-no loss of IHCs or OHCs in any condition (control: black; vehicle: red; ISRIB: blue). **C.** Number of synapses under control (black), vehicle (red), and ISRIB (blue) conditions is shown. Noise exposure led to synapse loss, which was mitigated by ISRIB. Pairwise 2-way ANOVA with repeated measures shows significant effect of treatment on paired (intact) as well as unpaired (orphan) synapses for control vs. vehicle (p = 0.0003 (paired)/0.004 (unpaired)) and vehicle vs. ISRIB (p = 0.0015/0.009), but not for control vs. ISRIB (p = 0.11/0.23). Data represent means ± s.e.

### Female CBA/J mice have higher baseline ABR wave-I amplitude than male mice

Whereas loss of IHC-SGN synapses is the characteristic anatomic finding of cochlear synaptopathy, the physiologic manifestation is reduction in the amplitude of wave I of the suprathreshold ABR, which serves as an indicator of the function and recruitment of SGNs stimulated by ribbon synapses onto IHCs^14^. Because sex differences in ABR wave-I amplitude have been observed in humans, we tested both male and female mice and adjusted for sex differences throughout all auditory physiology experiments^15^. First, we measured ABRs on a large cohort of 86 female and 98 male CBA/J mice between 6-8 weeks old. While there was no difference in ABR thresholds between male and female mice (**Figure 2A**), growth of wave-I amplitude with increasing stimulus presentation level and average suprathreshold (in response to 70-90 dB SPL stimuli) wave-I peak amplitudes were significantly higher in female mice between 8 and 40 kHz (p<0.0001, **Figure 2B-C**).

**Figure 2.**
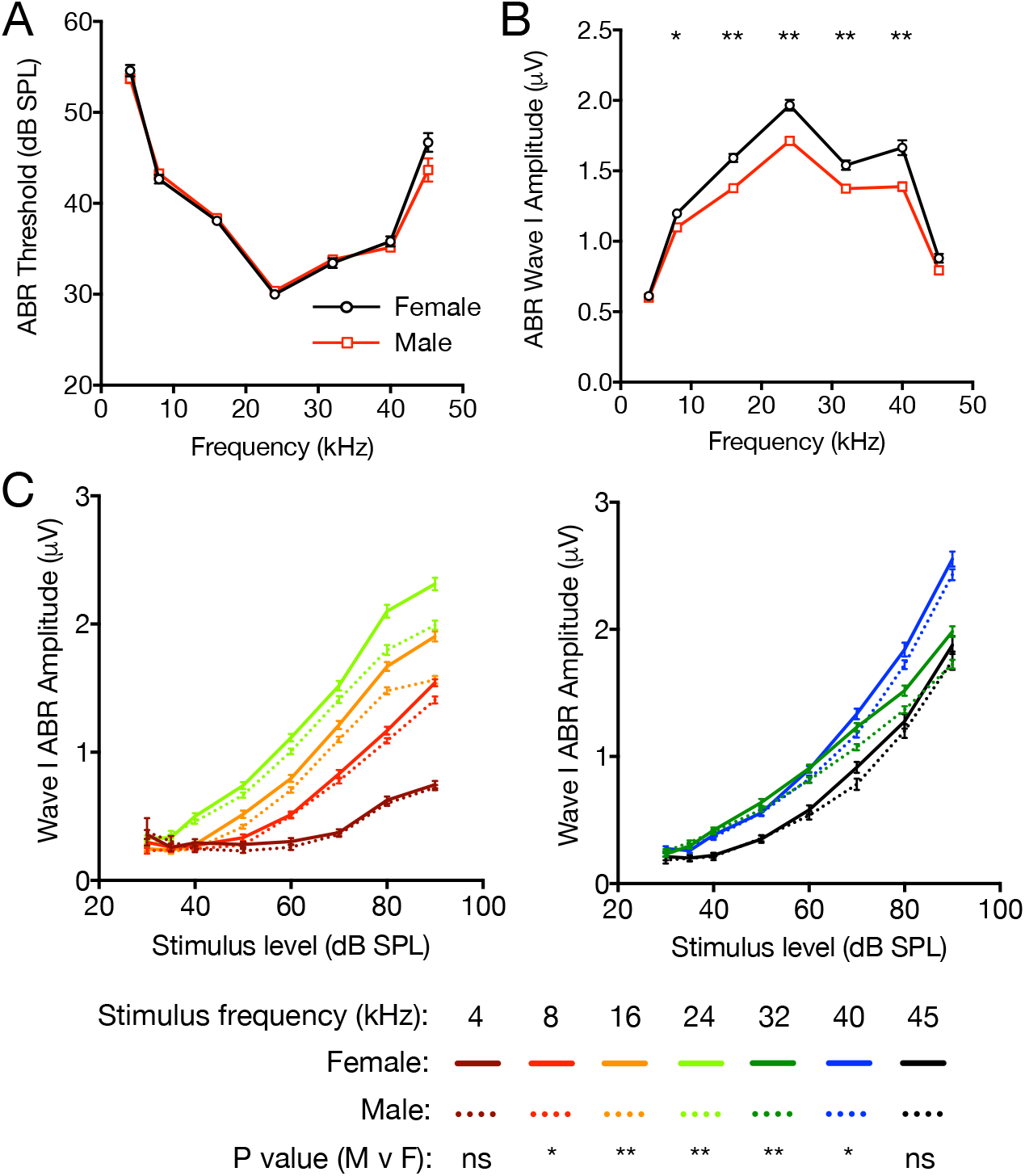
Baseline sex differences in ABR threshold and wave-I amplitude. 6-8-week-old female (black, N=86) and male (red, N=98) CBA/J mice underwent Auditory Brainstem Response (ABR) recording. ABR thresholds (**A**), average wave-I peak amplitudes (**B**) in response to suprathreshold tone pips at 70, 80 and 90 dB SPL, and growth of wave-I amplitude with stimulus presentation level (**C**) were measured. **A-B**: Values were compared between male and female mice at each frequency using Student’s unpaired two-tailed t-test with Bonferroni’s correction for multiple comparisons. **C**: Growth functions were compared at individual frequencies (low frequencies, left; and high frequencies, right) by 2-way ANOVA by sex and stimulus level. P values to evaluate statistical difference by sex, adjusted for stimulus level, are presented. Data represent means ± s.e. * p < 0.01; ** p < 0.001.

### Noise-induced ABR threshold and wave-I amplitude shifts differ in male and female mice

Having established these baseline sex differences, we sought to determine the efficacy of ISRIB to prevent cochlear synaptopathy across multiple noise-exposure levels in a sex selective manner. First, to precisely define our cochlear synaptopathy model, we exposed male and female mice to a range of sound pressure levels. 97.8 dB SPL 8-16-kHz octave-band noise produced immediate elevation in ABR thresholds at PNE1 that then returned to baseline by PNE21, accompanied by large permanent wave-I amplitude shifts (**Figure 3A-B, middle**). Lower noise-exposure levels produced similar outcomes— minimal threshold shift with more modest wave-I amplitude shifts (**Figure 3A-B, left**, for 94.3 dB SPL). Even slightly higher noise-exposure levels, however, gave rise to substantial permanent threshold shifts (**Figure 3A-B, right**, for 100.7 dB SPL; similar findings for 99.0 and 101.4 dB SPL, data not shown). As seen for baseline ABRs, the response to noise was sex-dependent: at higher noise-exposure levels (100.7 dB SPL), threshold shifts were larger in male mice, whereas at the lower noise-exposure levels (94.3 and 97.9 dB SPL), threshold shift was not sex-dependent, but noise-induced wave-I amplitude shift was significantly larger in female mice compared to males (**Figure 3C**, p < 0.05 for wave-I amplitude (bottom), but not threshold (top)). These findings led us to use only the 94.3 and 97.8-dB levels as a model in subsequent experiments for noise-induced cochlear synaptopathy without significant threshold shift.

**Figure 3.**
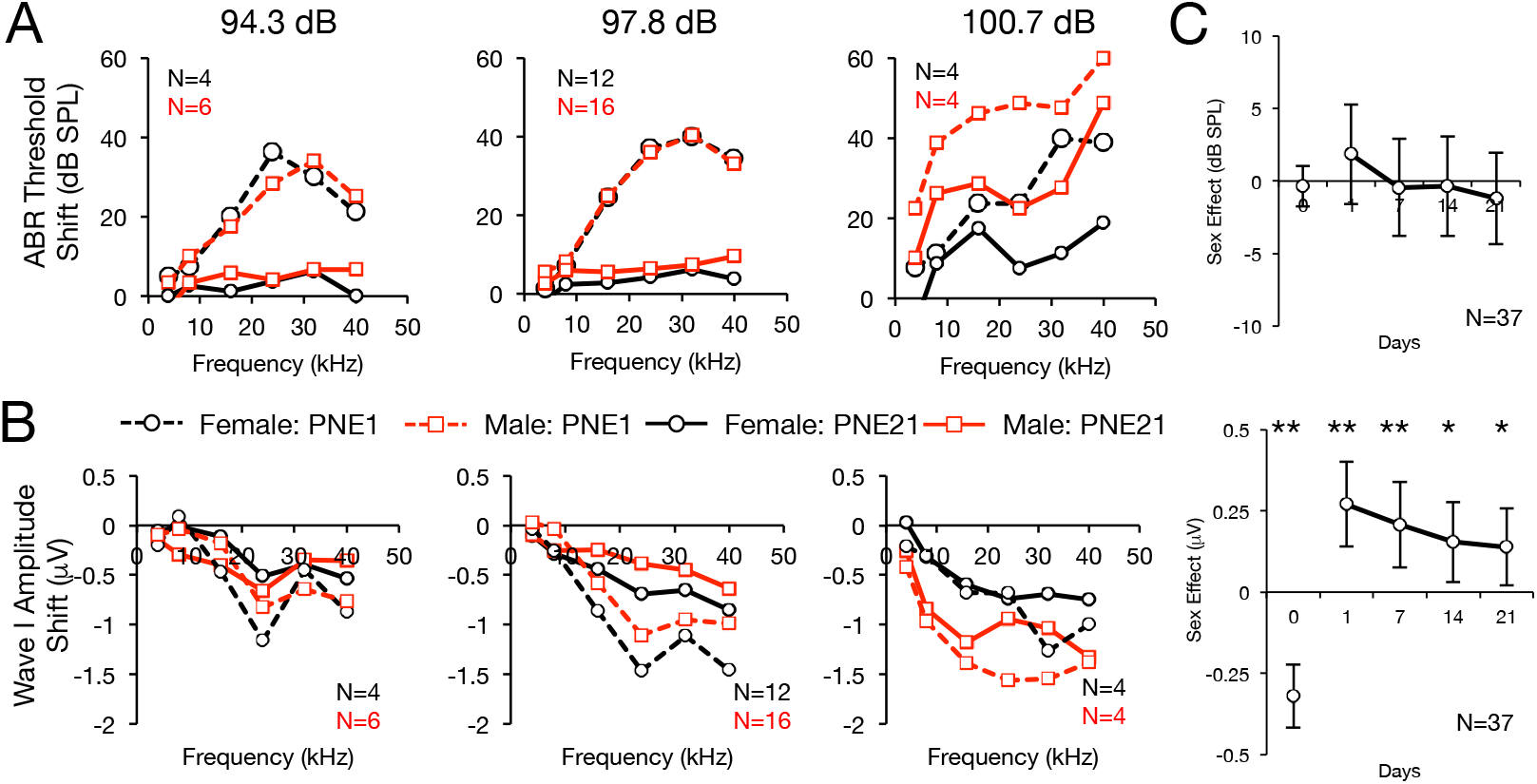
Sex differences in ABR threshold and suprathreshold wave-I amplitude shift after different levels of noise exposure. 8-week-old female (black) and male (red) CBA/J mice were exposed to 2h, 8-16 kHz octave-band noise at the indicated sound-pressure levels. ABRs were measured at baseline, 1 (dashed lines), and 21 (solid lines) days post noise exposure (PNE). Shifts in ABR thresholds (**A**) and suprathreshold wave-I amplitudes (**B**) relative to baseline were measured. 3-way ANOVA for sex, frequency, and noise exposure level indicated significant main effects of each factor on ABR threshold shift at PNE1, and on both threshold and wave-I shifts at PNE21, and interaction between sex and noise exposure level (p < 0.05 for all comparisons). Subgroup analysis at each noise-exposure level with 2-way ANOVA for sex and frequency revealed significant effect of sex at PNE21 on ABR threshold at 97.8 dB (p=0.02) and 100.7 dB (p=0.005), but not at 94.3, and on wave-I amplitude at 97.8 dB only (p = 0.02). **C.** Analysis was restricted to noise-exposure levels that caused cochlear synaptopathy without threshold shift (94.3 and 97.8 dB only). 3-way ANOVA reveals the effect of sex (male vs. female) on ABR threshold (top) and wave-I amplitude (bottom) at each timepoint, adjusted for noise exposure level and frequency. For baseline (Day 0), absolute ABR threshold and wave-I amplitude values were compared; for post-noise-exposure timepoints, ABR threshold and wave-I amplitude shifts (relative to baseline values) were compared. Positive effects indicate better hearing or less noise-induced hearing loss for male relative to female mice. Mean effect sizes ± 95% confidence intervals are shown. * p < 0.05; ** p < 0.01 for a significant effect of sex at each timepoint.

### Noise-induced ABR threshold and wave-I amplitude shifts are mitigated by ISRIB in male, but not female, mice

We then evaluated the effect of drug treatment (ISRIB vs. vehicle) in this cochlear synaptopathy model in male and female mice across the two noise-exposure levels (94.3 and 97.9 dB SPL) and multiple timepoints (0, 1, 7, 14, and 21 days after noise exposure). Threshold shifts were minimal for all mice compared to control, unexposed mice, though ISRIB-treated male mice did have slightly improved thresholds compared to vehicle-treated animals (**Figure 4A**). Wave-I amplitude shifts 1 day after noise exposure were lower in all ISRIB-treated animals. By 21 days after noise exposure, however, only male mice retained the protective effect of ISRIB (**Figure 4B**). This is in contrast to noise-induced synapse reduction, which was mitigated by ISRIB in male and female mice equally (**Figure 5**). Direct comparison of synapse counts and wave-I amplitudes in ISRIB-treated male and female mice demonstrated significantly higher wave-I amplitudes for equivalent synapse counts in male compared with female mice (**Figure 5C** (right), p < 0.05).

**Figure 4.**
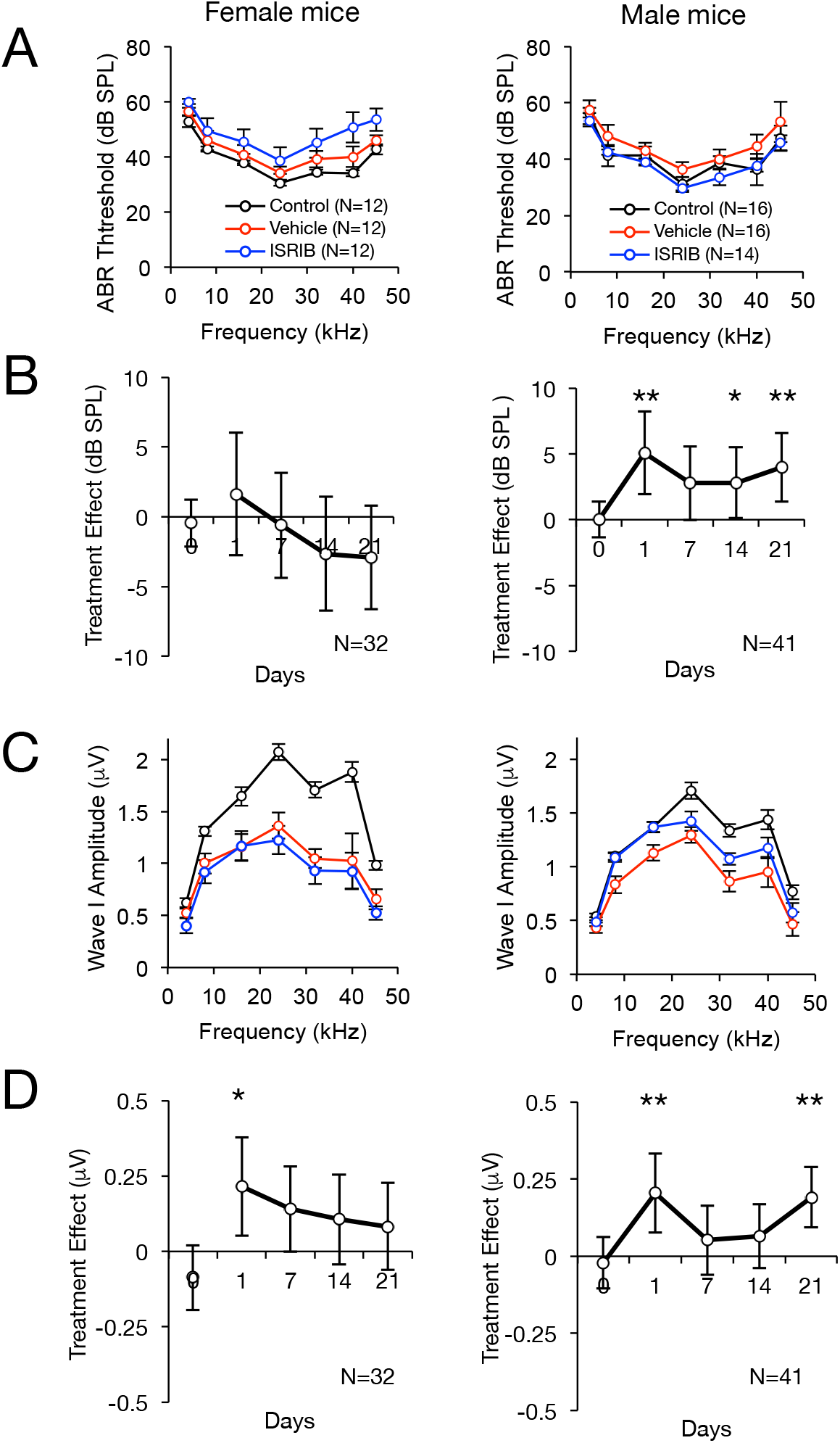
Effect of ISRIB on treatment of ABR threshold and suprathreshold wave-I amplitude in noise-induced cochlear synaptopathy. ABR threshold (**A,B**) and suprathreshold Wave-I amplitudes (**C,D**) were measured in female (F, left) and male (M, mice) CBA/J mice after a single intraperitoneal dose of ISRIB (ISR) or vehicle (VEH) and exposure to 8-16 kHz octave-band noise. **A,C** Unadjusted ABR measurements in mice 21 days after 97.8 dB SPL noise exposure demonstrated no significant differences in ABR threshold (**A**) between control (no noise exposure, black), vehicle- (red), and ISRIB-pretreated (blue) mice, but significant reduction in wave-I amplitude (**C**) in both vehicle- and ISRIB-treated female mice, with partial protection with ISRIB in male mice. **B,D.** In an adjusted timecourse analysis including mice exposed to 94.3 dB SPL (4 F/VEH, 4 F/ISR, 5 M/VEH, 6 M/ISR) and 97.8 dB SPL (12 F/sVEH, 12 F/ISR, 16 M/VEH, 14 M/ISR) noise, 3-way ANOVA (for treatment, noise-exposure level, and frequency) revealed main effects of all three factors without interaction effects (p < 0.05 for all). Multiple linear regression was used to quantify the effect of drug treatment (ISRIB vs. vehicle) in female (left) and male (right) mice at each timepoint, adjusted for noise exposure level and frequency. For baseline, absolute ABR threshold (**B**) and wave-I amplitude (**D**) values were compared; for post-noise-exposure timepoints, ABR threshold (**B**) and wave-I amplitude shifts (**D**) (relative to baseline values) were compared. Positive effects indicate better hearing or less noise-induced hearing loss for the ISRIB relative to Vehicle condition in all graphs. Mean effect sizes ± 95% confidence intervals are shown. * p < 0.05; ** p < 0.01 for a significant effect of ISRIB treatment at each timepoint.

**Figure 5.**
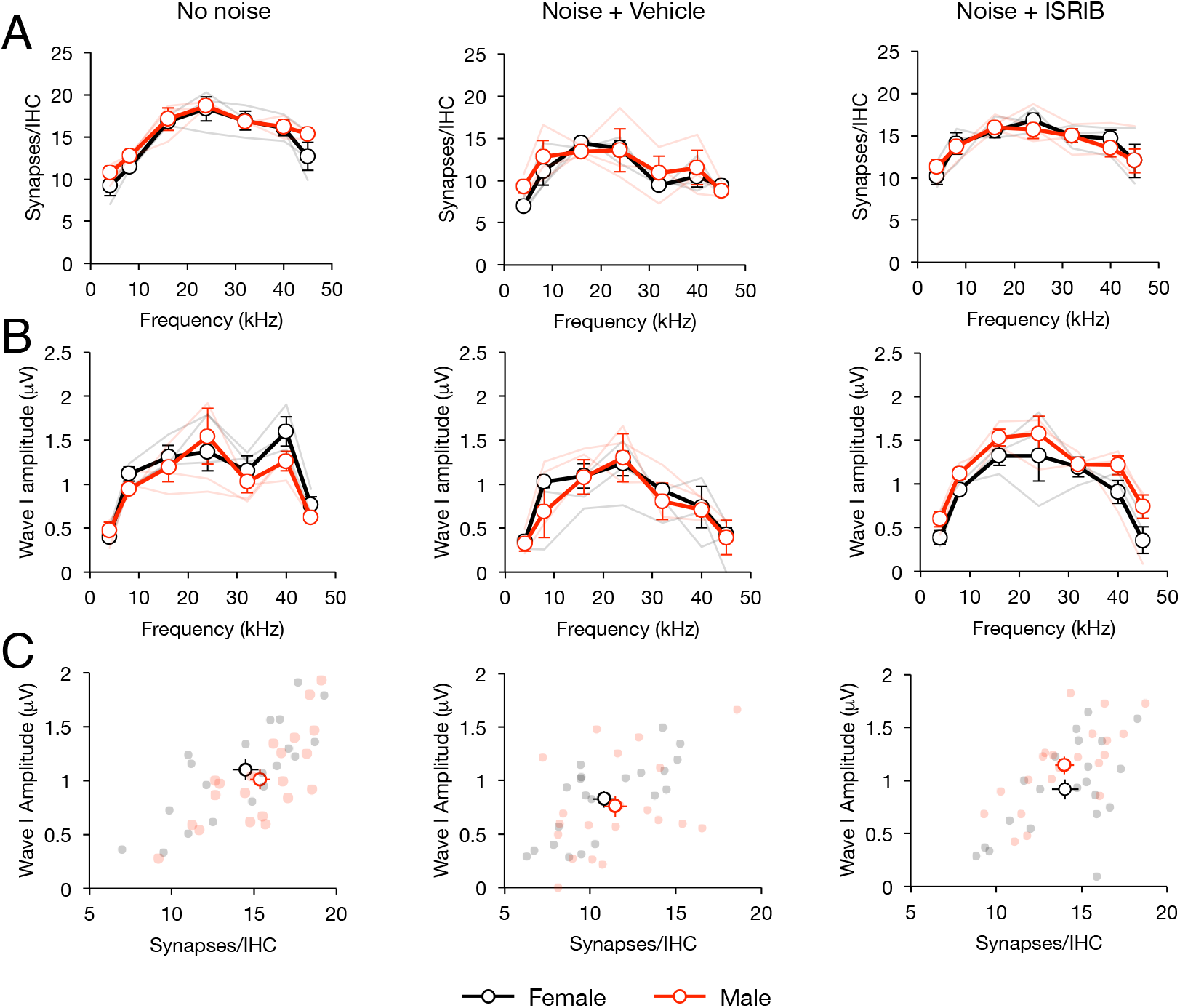
Individual-level comparison of treatment and sex effects on noise-induced synapse loss and wave-I amplitude decrease. Female and male mice were pre-treated with ISRIB or vehicle and exposed to 97.8 dB SPL noise, or unexposed (control; N=3 in each sex/treatment condition). ABR was performed at PNE21 and synapses quantified at PNE45. **A.** Comparison between female (black) and male (red) mice reveal no significant sex difference in synapse counts in control (left), vehicle (middle), or ISRIB (right) conditions. **B**. Female and male control (left) and noise + vehicle-treated (middle) mice had similar suprathreshold wave-I amplitudes, but noise + ISRIB-treated male mice had significantly improved wave-I amplitudes compared to female mice (right, p = 0.0036, 2-way ANOVA). **C.** Wave-I amplitude and synapse correlation plots demonstrate no significant differences between male and female mice in control (left) and noise + vehicle (middle) conditions, but significant shift towards higher wave-I amplitude for similar synapse number in male, compared to female, mice under noise + ISRIB conditions (p = 0.028, t-test). Data represent means ± s.e., with individual animals shown in shaded lines or dots.

## Discussion

Our anatomic and physiologic findings demonstrate that ISRIB treatment can reduce cochlear synaptopathy and “hidden hearing loss,” an emerging phenomenon that may underlie functional auditory disability for millions of individuals. This finding extends our previous work demonstrating the efficacy of ISRIB in preventing permanent noise-induced auditory threshold shifts into the related phenomenon of cochlear synaptopathy, and suggests that a common, targetable mechanism — activation of the UPR — may underlie noise-induced hair-cell degeneration and synapse loss in the cochlea^4^. Additionally, this work expands into sensory systems a large body of research findings in neurological disease models demonstrating the ability to target the UPR to prevent or reverse synaptopathy. Our current finding that ISRIB protects against synaptopathy in the cochlea, therefore, strongly corroborates previous findings in the brain. A link between UPR-mediated neurodegeneration in the brain, hair-cell loss, and cochlear synaptopathy was suggested by recent work in mice deficient for the ER-stress-regulator mesencephalic astrocyte-derived neurotrophic factor (MANF). In this genetic mouse model, ER stress is associated with loss of synapses in the cochlea, accompanied by progressive hair-cell death and hearing loss^16^. The findings that MANF is cytoprotective in neurons and that *Manf* inactivation is associated with loss of synapses in the inner ear suggests that the UPR may act similarly in the brain and auditory sensory cells. Our current findings are consistent with this genetic model. However, our work is the first to provide direct evidence of UPR involvement in an acquired, noise-induced model of cochlear synaptopathy in wild-type mice.

The mechanism by which noise exposure may act through the UPR to cause cochlear synaptopathy is not known, but insight can be obtained through previous work in the central nervous system that may be relevant to the cochlea’s response to noise. Models of mechanical damage, such as occurs in NIHL, have implicated the UPR in neurons. In a study of crushed optic nerves, treatment of mice with ISRIB reduced axonal degeneration^17^, and UPR modulation improved recovery after spinal cord injury in mice^18^. Our work in genetic and noise-induced mouse models has similarly demonstrated a role for ER Ca^2+^ dysregulation as a link between acoustic overstimulation and UPR activation^4^. Cochlear supporting cells exhibit periodic ER Ca^2+^ release upon acoustic overstimulation^19^, and mice deficient in Tmtc4, a novel deafness gene, exhibit impaired ER Ca^2+^ reuptake, elevated UPR gene expression, and profound post-natal progressive hearing loss^4^. It is possible that noise-induced disruption of ER Ca^2+^ dynamics may affect synaptic function; the ER is the primary store of intracellular Ca^2+^ in presynaptic terminals, and Ca^2+^ is a critical regulator of neurotransmitter release, hair-cell vesicle recruitment, and regulation and timing of vesicle fusion in ribbon synapses.^20^ UPR-related modulation of Ca^2+^ homeostasis alters synaptic transmission and plasticity, and ribbon synapse physiology depends on intracellular Ca^2+^ buffering^21^. Inhibitors of ER Ca^2+^ uptake, including thapsigargin, a known UPR inducer, increase resting cytoplasmic Ca^2+^ levels and delay recovery time after depolarization in IHCs^22^. Finally, ryanodine, which stimulates ER Ca^2+^ release, leads to the reversible inhibition of compound action potentials in the auditory nerve, presenting further support of the role of Ca^2+^ in hair-cell synaptic transmission.^23^

In the current study, we also show that the protective effect of ISRIB, as well as the basic phenomenon of noise-induced cochlear synaptopathy, is highly sex-dependent. Female mice had higher ABR wave-I amplitudes at baseline (**Figure 2**), were more susceptible to noise-induced reduction in suprathreshold amplitudes (**Figure 3**), and were less responsive to ISRIB treatment compared to male mice (**Figure 4**). These findings are similar to published reports in humans, where women exhibit larger baseline wave-I amplitudes despite similar auditory thresholds, as well as greater sensitivity to wave-I amplitude decrease, when compared to men^15,24^. These physiologic sex differences were not reflected, however, in baseline, noise-evoked, or ISRIB-treated synapse anatomy (**Figures 1 and 5**); the mechanisms underlying this discordance between synapse anatomy and function require further investigation. More generally, our study highlights the critical importance of including sex as a biological variable in the investigation of treatments for NIHL and cochlear synaptopathy ^25^, as well as in the evaluation of UPR modulators such as ISRIB in all disease models.

In conclusion, we have shown that targeting the UPR with ISRIB is an effective means of preventing cochlear synaptopathy in this mouse model, and that with careful and appropriately powered studies, we were able to determine that these effects are sex-dependent. We have previously demonstrated that the UPR is involved in the pathogenesis of permanent NIHL, and that ISRIB can prevent hair-cell death^4^, which is otherwise irreversible. In cochlear synaptopathy, hair cells do not die, but synapses are lost. Unlike hair-cell regeneration, synaptic regeneration in the cochlea does occur^26^; the ability of ISRIB and other UPR modulators to promote synaptic plasticity and regeneration thus make the UPR an extremely appealing target pathway to treat multiple modes of noise-induced hearing loss — both permanent threshold shift, through prevention of apoptosis and hair-cell death, and cochlear synaptopathy, through prevention of synapse loss and promotion of synaptic regeneration.

## Methods

### Auditory testing and noise exposure

ABR thresholds and intensity functions (10-90 dB SPL) were recorded in response to 4-kHz tone pips in mice at baseline and 1, 7, 14, and 21 days post noise exposure (PNE) as described (RZ6, Tucker-Davis Technologies)^4^. 8-week-old mice were exposed to 94.3, 97.8, 99.0, 100.7, or 101.4 dB SPL 8-16 kHz octave-band white noise for 120 minutes in a custom-built, calibrated, reverberant sound chamber^24^. Animals were awake and unrestrained in an acoustically transparent wire cage on rotating platform throughout noise exposure. ABR thresholds were determined as described^4^ and wave-I amplitude measured as the difference between wave-I peak and trough by an investigator blinded to treatment group and confirmed with a custom peak detector (MATLAB). Suprathreshold ABR wave-I amplitude was defined as the average wave-I amplitude in response to stimuli at 70, 80, and 90 dB SPL. To assess the effect of UPR modulation on NIHL, animals were pretreated with intraperitoneal injection of 2.5 mg/kg ISRIB (MilliporeSigma) or its vehicle (50% DMSO, 50% PEG-400) 2 hours prior to noise exposure.

### Experimental Rigor and Statistical Analysis

Wild-type male and female CBA/J mice were obtained from Jackson Laboratories at 6-7 weeks and allowed to acclimatize to our facility before any procedures. Baseline auditory testing was conducted, and mice were randomly assorted into treatment and noise exposure groups for noise exposure at 8 weeks of age. Noise exposures were performed on four mice at a time, one from each treatment group: male and female treated with ISRIB or vehicle. All ABR measurements were performed and scored by an independent investigator blinded to treatment group.

Specifics of statistical analysis are described in figure legends. Pairwise comparisons were performed with two-tailed, unpaired Student’s *t*-test. 2- or 3-way ANOVA was used to assess statistical significance of differences attributable to sex, treatment, noise exposure level, and timepoint with subgroup analyses and correction for repeated measures when appropriate. Multiple linear regressions were used to quantify the effect of drug treatment and sex effect. A p value of <0.05 was considered significant. Tests were performed with MATLAB and GraphPad.

### Immunohistochemistry and Synapse Quantification

Synapse quantification was performed by examining co-localization of immunohistochemical staining for CtBP2 and GulA2. Cochleae were perfused with 4% paraformaldehyde, removed from temporal bones, and fixed for 2 hours at room temperature. Cochleae were then decalcified in EDTA at room temperature for 5 days, dissected, and blocked in 5% normal horse serum in PBS with 0.3% Triton X-100 for one hour at room temperature. Primary antibodies were diluted in blocking solution of 1% normal horse serum in PBS with 0.3% Triton X-100 and incubated overnight at 37°C as follows: mouse (IgG1) anti-CtBP2 (C-terminal Binding Protein) 1:200 (#612044, BD Transduction Labs) targeting pre-synaptic ribbons; mouse (IgG2a) anti-GluA2 (Glutamate receptor subunit A2) at 1:2000 (#MAB397, Millipore) for labeling post-synaptic receptor patches; and rabbit anti-Myosin7a at 1:200 (#25-6790 Proteus Biosciences) for delineating hair cells. Species-specific secondary antibodies were applied for two 60 minute incubations at 37°C as follows: Alexa Fluor 488-conjugated goat anti-mouse (IgG2a) at 1:1000 (#A21131, Life Technologies); Alexa Fluor 568-conjugated goat anti-mouse (IgG1) at 1:1000 (#A21124, Life Technologies); Alexa Fluor 647-conjugated chicken anti-rabbit at 1:200 (#A21443, Life Technologies). Cochleae were mounted using Vectashield mounting medium (Vector Labs).

A confocal microscope (Nikon Ti) with a 60x oil-immersion objective (1.4 NA) was used to acquire z-stack images, with 0.25 μm step size, of the entire synaptic region of IHCs. Image stacks were imported to Imaris processing software to be analyzed in three dimensions. Individual IHCs were quantified by an investigator blinded to treatment, sex, and cochlear frequency. Pre-synaptic ribbons, glutamate receptors, and colocalized intact synapses were counted using stain from Myosin7a as well as nuclear stain from anti-CtBP2 to delineate individual cells.

Cochleograms were generated from 10x images of all pieces of cochlea. Cochlear lengths were mapped in Imaris and frequencies calculated based on strain-specific data^27^ Missing hair cells were counted from anti-Myosin7 immunostain in 10x images collected to generate the cochleogram. Hair cells were counted in segments of 200 μm centered around each frequency and expressed as percentage of missing hair cells.

### Study Approval

The UCSF Institutional Animal Care and Use Committee approved the animal studies.

## Acknowledgements

This work was supported in part by a grant from the National Institute on Deafness and Other Communication Disorders (NIDCD) to DKC (R03DC015082), from Hearing Research, Inc., to DKC, and from the National Institute of Neurological Disorders and Stroke (NINDS) to EHS (2R01NS058721)

## Author Contributions

SLR, EHS, and DKC created the initial design. DKC and EHS oversaw experiments and ethics. SLR and IRM contributed to the experiments. SLR, IRM, JL, and DKC analyzed and interpreted data. SLR, JL, EHS, and DKC drafted the manuscript.

## Additional Information

### Competing Interest Declaration

Drs. Sherr and Chan are founders and shareholders in Jacaranda Biosciences. Dr. Sherr is listed as an inventor on the published patent application: Novel methods of treating hearing loss (PCT/US2016/058348) Authors Rouse, Li, Matthews declare no competing interests.

